# Domestic gardens as favorable pollinator habitats in impervious landscapes

**DOI:** 10.1101/374116

**Authors:** Marine Levé, Emmanuelle Baudry, Carmen Bessa-Gomes

## Abstract

Urban expansion is correlated to negative biodiversity trends. The amount of impervious surfaces in urban areas is a determinant of pollinator species assemblages. While the increase in urbanization and impervious surfaces negatively impacts pollinators, cities also encompass urban green spaces, which have a significant capacity to support biodiversity. Among them, domestic gardens that represent a non-negligible fraction of green spaces have been shown to benefit pollinators. Domestic gardens may form habitat clusters in residential areas, although their value at a landscape scale is still unknown. Here, we investigate the combined effects of impervious surfaces and domestic garden areas on pollinator richness. Due to the difficulty of accessing privately owned domestic gardens, we chose to use citizen science data from a well-established French citizen science program known as SPIPOLL. Using regression tree analysis on buffers located from 50m to 1000m around the data points, we show the importance of pollinators being in close proximity to domestic gardens as locally favorable habitats that are embedded within a landscape, in which impervious surfaces represent unfavorable areas. We highlight the inter-connection between local and landscape scales, the potential for patches of domestic gardens in residential areas, and the need to consider the potential of gardeners’ coordinated management decisions within a landscape context.

**Highlights:**

- Citizen science provided access to domestic gardens, understudied urban green spaces
- Impervious surfaces limit pollinators presence at landscape level
- Sufficient critical amount of gardens increased pollinator diversity at local scale
- Critical amount of gardens’ knowledge may favor coordinated decisions by gardeners
- Pollinators may benefit from patches of domestic gardens in an urban matrix

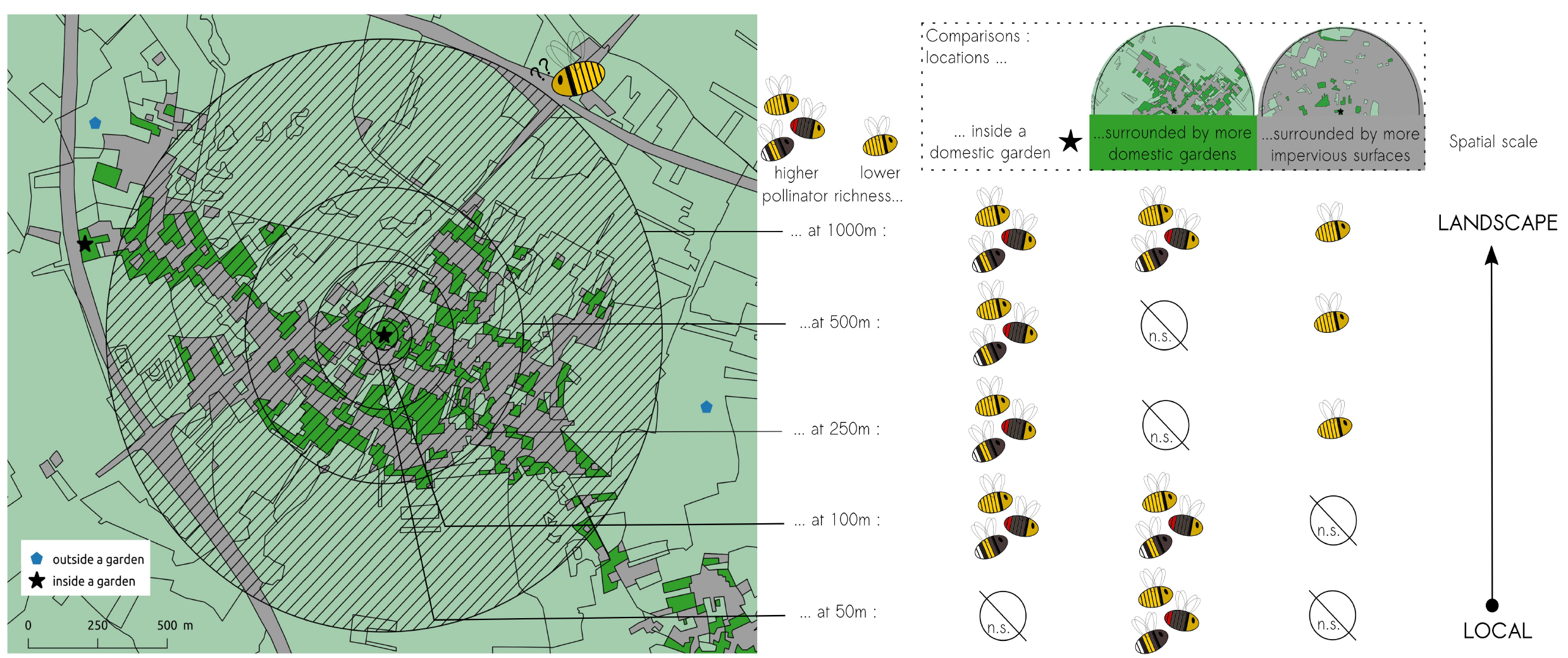

## 1 Introduction

Urban areas now contain more than half of the world’s population and will continue to grow (United Nations, 2014). As urban expansion leads to an increasing number of former natural and semi-natural areas becoming urbanized, urbanization is now considered to be a major threat to biodiversity (Grimm et al., 2008). General findings stress the negative impact of urban areas on biodiversity through habitat loss, reduced habitat quality, and habitat homogenization (McKinney, 2008), as well as their impact on native species extinction (Czech et al., 2000). The increasing proportion of impervious surfaces in urban areas is a possible proxy for urbanization level. It is also a major determinant for several species assemblages, such as bees (Fortel et al., 2014; Geslin et al., 2016) and amphibians (Parris, 2006), as well as for urban ecosystem functions, such as water resources and flow regulation (Arnold and Gibbons, 1996).

Cities encompass urban green spaces (UGS), such as cemeteries, urban wasteland, public gardens, community gardens, and domestic gardens (Lepczyk et al., 2017). These UGS account for a variable percentage of a city’s area, ranging from 2 to 46% in European cities (Fuller and Gaston, 2009). Their ability to support biodiversity has been recently acknowledged (Aronson et al., 2014; Beninde et al., 2015), and there is now a call to effectively integrate UGS in biodiversity planning and management to ensure their full inclusion in biodiversity conservation (Lepczyk et al., 2017). They may constitute a diversification of land usages given the general impervious surface and thus support increased levels of biodiversity (McKinney, 2008). Moreover, UGS benefit human health and well-being (Tzoulas et al., 2007).

Domestic gardens are an understudied type of UGS, mainly because of their limited accessibility to researchers and their supposed non-relevance to conservation (Cameron et al., 2012; Cook et al., 2012; Goddard et al., 2010). Yet domestic gardens may account for a large part of UGS and are thus worth considering in terms of their contribution to biodiversity conservation. Their estimated areas in cities vary from 16% in Stockholm, Sweden (Colding, 2007) to 22-27% in the UK (Loram et al., 2007) and 36% in Dunedin, New Zealand (Mathieu et al., 2007). Their distribution is heterogeneous within cities and surrounding regions: in Flanders, there is a lower concentration of gardens in city centers, but a higher proportion in the areas surrounding the centers and peri-urban areas (Dewaelheyns et al., 2014). Various organisms have been found to benefit from urban or peri-urban domestic gardens (Goddard et al., 2010), such as birds (Daniels and Kirkpatrick, 2006; van Heezik et al., 2008) and invertebrates (Smith et al., 2006a, 2006b; Sperling and Lortie, 2010), including pollinators (Pardee and Philpott, 2014).

In this study, we chose to focus on pollinators because of their role in ecosystem functioning (e.g. Potts et al., 2010 but also Kleijn et al., 2015), but also because of their adaptation to urban environments and the challenge associated with the low mobility of many small solitary bee species (Greenleaf et al., 2007; Zurbuchen et al., 2010). We thus consider pollinator richness as a surrogate for domestic gardens biodiversity. Indeed, grouped domestic gardens, i.e. patches, may represent more favorable habitats to pollinators with small flight ranges, as they are able to take advantage of the nearby resources, either inside the garden or in adjacent gardens (Lerman et al., 2018). As urban perturbation is particularly high on small spatial scales and eliminates close living species (McKinney, 2008), suitable habitat patches such as domestic gardens could serve as refuges for pollinators. Hinners et al. (2012) found that resources are insufficient to maintain high pollinator diversity in suburban habitats less than 80,000m^2^, while in habitats around 200,000m^2^, richness was comparable to semi-natural areas: a threshold value for pollinator conservation might lie between these two figures. However, these areas are already considerably greater than the average domestic garden size, estimated to be 571m^2^ in Belgium (Dewaelheyns et al., 2014) and 190m^2^ in the UK (Davies et al., 2009). Yet the peripheries of many Western cities consist of extended suburban areas comprising residential areas with detached houses and private gardens. Consequently, the combined surface of neighboring domestic gardens might attain the threshold value of habitat patch size. When identifying actions to reduce the impact of urbanization on pollinators, the attained threshold surface may be an indicator of their efficiency.

In this context, pollinators are an interesting choice when studying domestic gardens as their decline is made visible to citizens, and individual actions in favor of pollinators are accessible to garden owners. While urbanization adversely affects pollinators by destroying floral resources and nesting sites (McKinney, 2002), the installation of “bee hotels” (artificial structures with materials that bees can use as a nesting site, such as wooden blocks with holes, paper tubes, etc.) has a variable impact on pollinator richness and abundance depending on the pollinator species (Gaston et al., 2005; MacIvor and Packer, 2015). The planting of pollinator-friendly flowers likewise has a variable impact depending on the chosen flower species and targeted insect species (Garbuzov and Ratnieks, 2014; Salisbury et al., 2015). A more precise determination of the scale of effect of domestic garden patches on pollinator richness in peri-urban areas would make an important contribution to biodiversity planning and management.

Because of their privately owned status, domestic gardens are not easily accessible and are thus less often the subject of research compared to other types of UGS (Hernandez et al., 2009). While obtaining regular access to gardens is difficult, long-term data gathering from domestic gardens is still possible through citizen science programs. The French citizen science program known as SPIPOLL was launched in 2010 (Deguines et al., 2012) by the National Museum of Natural History (MNHN) and Office for Insects and their Environment (OPIE) with a focus on flower-visiting insects, most of which are insect pollinators. Using a short protocol, SPIPOLL allows participants to take photographs of insects seen on flowers and send them to an internet database. The collected photographs result in an understanding of insects and their land-use preferences (Deguines et al., 2012).

SPIPOLL is a nation-wide program, although we choose to focus on the Île-de-France region in this study. Île-de-France is a densely populated region and is representative of urban areas in Western industrialized countries with their organization around a metropolis. The Parisian metropolis is located approximately in the center of the Île-de-France region and is surrounded by successive urban belts with decreasing urbanization, with a higher urban concentration around transportation networks (Fig. 1). Semi-detached or detached houses surrounded by domestic gardens are more frequent on the Paris periphery. Overall, the region allows us to study an urbanization gradient with a variable proportion of built-up, residential, and garden areas.

**Figure 1.**
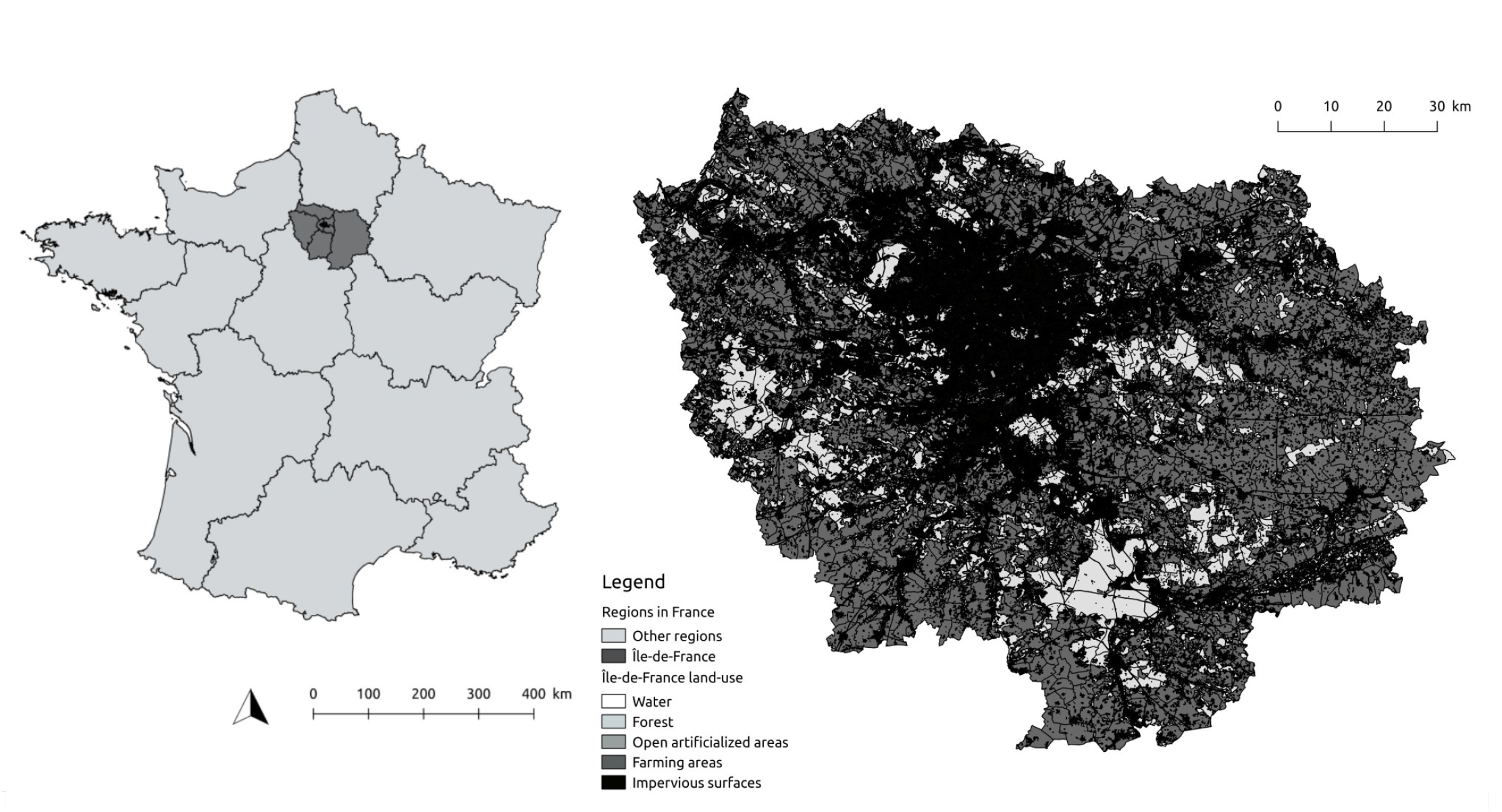
Location of the studied region in France and land use in Île-de-France.

Determining a threshold for cumulative domestic garden areas that benefit pollinators relative to urban impervious surfaces on a given scale could help urban planners in the decision-making process. Moreover, regarding citizen science programs such as SPIPOLL, participants’ knowledge of this threshold and the geographic situation of their garden could help them to better appreciate pollinator diversity as well as local pressures, and thus consider this diversity relative to the local urbanization stage and processes, especially as the inhabitants may also experience these to some extent.

The present study aims to understand the effect of urbanization and domestic gardens on pollinator richness and their relative importance. It includes several spatial scales relevant to the flight distances of pollinators and the size of domestic garden patches in peri-urban residential areas. Our hypotheses are as follows: (1) the effects of gardens on pollinator richness will be limited in densely urbanized areas, in which domestic gardens may not be determinants of pollinator richness; and (2) in areas where domestic gardens do have an influence on pollinator richness, the latter will be higher in areas with a low proportion of impervious surfaces and a high proportion of gardens.

## 2 Methods

The use of a French citizen science program focusing on pollinators allowed us to gather a large amount of data from locations that are usually difficult to access, i.e. domestic gardens. As we used citizen science data, several factors out of our control and irrelevant to this study may be related to data variability, such as temperature and cloud coverage (see Data selection method section below). Likewise, data location depended on the participants’ decision to contribute to the program and resulted in spatial autocorrelation. To account for these factors, we used a two-step methodology to answer our research question. We initially standardized pollinator data via a first modeling step (see Data standardization method section below). We then used the residuals of this first modeling step to analyze the dependency of pollinators on the variables of interest: impervious surface proportion, domestic garden surface proportion, and location of data points inside the gardens (see Regression tree modeling method section below).

### 2.1 Diversity of flower-visiting insect data and standardization

#### 2.1.1. Data selection

The SPIPOLL program started in 2010 and has already led to several publications (e.g. Deguines et al., 2012; Deguines et al., 2016; Desaegher et al., 2018). The SPIPOLL protocol is as follows: participants have to take photographs of insects visiting a given flower species over a period of at least 20 min. They then identify the insects using an online key based mostly on morphological features and upload one photograph per identified insect to the dedicated website (www.spipoll.org). Identification is conducted at the level of morphotypes, representing groups of individuals at six various taxonomic levels: a species, a species from a genus, a genus, a species from different genera, several genera within a family, and a whole family. SPIPOLL encompasses 630 different morphotypes, of which 95.4% are considered to be pollinators. Thus, even if the program takes into account all flower-visiting insects, it mostly focuses on pollinators, a term that we will subsequently use. A set of all photographs taken during the same session is called a collection. Collections are spatially located by participants when they upload the photographs, and the flower species of the collection, or at least the flower family, is recorded. Because of the design of the SPIPOLL program, collections are considered to be units of observation. They will therefore be used as the unit for richness analysis in this study. Insect identifications are verified by several professional entomologists (see Deguines et al., 2016). Additional information is also gathered: wind intensity on a five-point scale from no wind to strong and continuous wind, temperature in four categories (<10°C, 10-20°C, 20-30°C, >30°C), sky cloud coverage in four categories (0-25%, 25-50%, 50-75%, 75-100%), and presence of shade on the observed flower (yes/no). A full description of the sampling protocol may be found in Deguines et al. (2012).

We obtained all data points recorded in Île-de-France from 2010 to 31 October 2017. We selected data points with wind intensity belonging to the first four of five categories, temperatures of 10- 20°C, 20-30°C, and >30°C, and no shade on the flower. We excluded data points with an observation period of less than 20 min. We only retained data points for which the flower belonged to a family appearing more than 30 times in the dataset (Fig. 2). We also verified for suspicious data points, i.e. those with a particularly high number of insects recorded compared to the observation time. We finally obtained 2470 data points.

**Figure 2.**
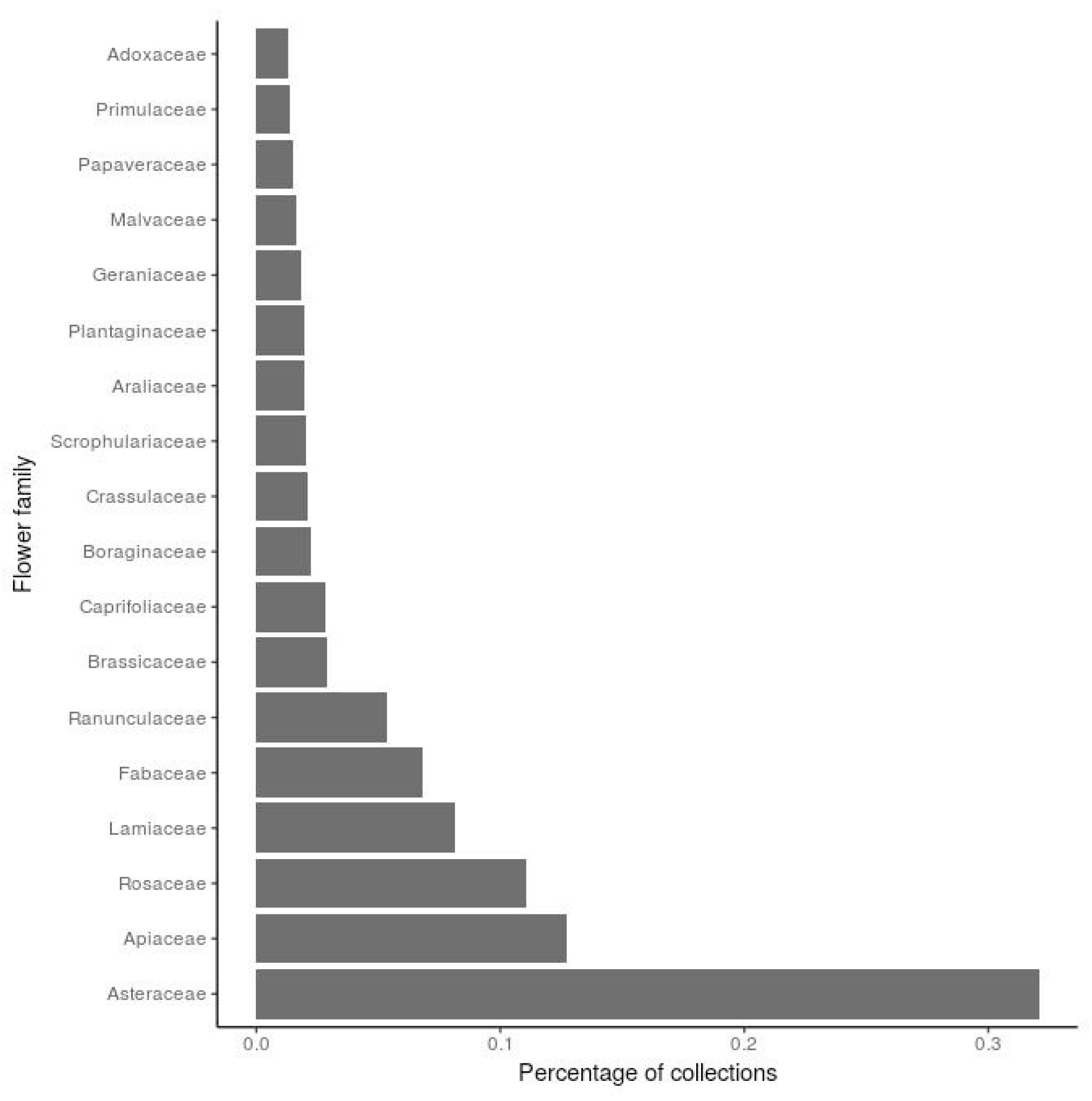
Flower families which are present in 30 recorded collections and more, ordered by the number of collections in which they are present.

SPIPOLL participants are free to observe pollinators anywhere, not only in their gardens. We thus identified data points recorded in domestic gardens using a Geographic Information System map reporting land use categories over the Île-de-France region, the MOS database (Institut d’Aménagement et d’Urbanisme, 2012). Data points located within the MOS category of “individual habitat gardens” were labeled as “inside gardens” and the remaining as “outside gardens,” creating a dichotomous garden local context variable (1/0). We also included data points located in a 20m buffer zone around the MOS category of “individual habitat gardens,” as participants have to locate the session on a map, which may lead to inaccurate locations if a large-scale view is used.

As species identification was conducted at a morphological level, we calculated the morphotype richness for each data point and standardized it per 20 min in order to account for variable observation duration.

#### 2.1.2. Data standardization

We observed spatial autocorrelation within data (see Supplementary Fig. S1), probably due to participants’ tendency to record collections around their habitation. As collections recorded near an inhabitant’s garden may have had quite a different context compared to collections recorded inside a garden, we retained all data points instead of excluding closely located data points. In order to manage autocorrelation, we built a generalized least square (GLS) model using the log of morphotype richness per 20 min as the dependent variable, because of its skewed distribution. We included latitude and longitude in an exponential spatial autocorrelation structure and added weather variables (i.e. temperature, cloud coverage, wind intensity), temporal variables (i.e. hour, week, year), flower family, and observer identity. As one aim of this study was to disentangle the local effect of the collection location in a garden from the larger effects of land use around the collection, we also used the dichotomous garden local context variable to indicate whether collections were located inside (1) or outside (0) a garden. We built one GLS model including this garden local context variable and another excluding it, hereafter respectively referred to as the models standardized and unstandardized for local context. Latitude, longitude, week, and hour were numerical variables. Cloud coverage, wind intensity, temperature, flower family, year, and garden were categorical variables. We assumed that all these variables had a linear effect, while we added a quadratic effect for hour and week based on the known seasonal and diurnal patterns of pollinators (Herrera, 1990; Knop et al., 2017; Lefebvre et al., 2018). We initialized spatial autocorrelation structure parameters after preliminary analysis of the data semi-variogram. We chose values that could capture short range effects: 400m for range effect, i.e. accounting for a leverage of autocorrelation effect with distance; and 0.9 for nugget effect, i.e. accounting for data which structure has a shorter range than sampling interval. We used the GLS function provided by the nlme package for R (Pinheiro et al., 2017). Residual spatial autocorrelation was verified using Moran’s I and along 100m steps for 20 intervals, which covered the full range of the chosen environmental buffers.

### 2.2 Landscape structure and composition

We used a geographic information system (GIS) to treat land use data. We also used the MOS database for general land use in Île-de-France (IAU). MOS covers the Île-de-France region with a 25m resolution and is regularly updated every 4-5 years. The present study uses the last updated version from 2012. MOS categories include 81 items in the most precise version; we used a version with 15 categories.

We extracted permeable areas and built impervious surfaces using Quantum-GIS (QGIS Development Team, 2016) in several buffers around the collections’ data points. Buffer sizes were chosen based on the literature on wild bee flight distances (Zurbuchen et al., 2010): 0m, 100m, 250m, 500m, and 1000m. Forests, artificial open spaces, water, farming areas, semi-natural areas, and domestic gardens were classified as permeable surfaces. Transport networks, facility areas, collective and individual housing areas, activity areas, building sites, and landfills were classified as built impervious surfaces (Fig. 3).

**Figure 3.**
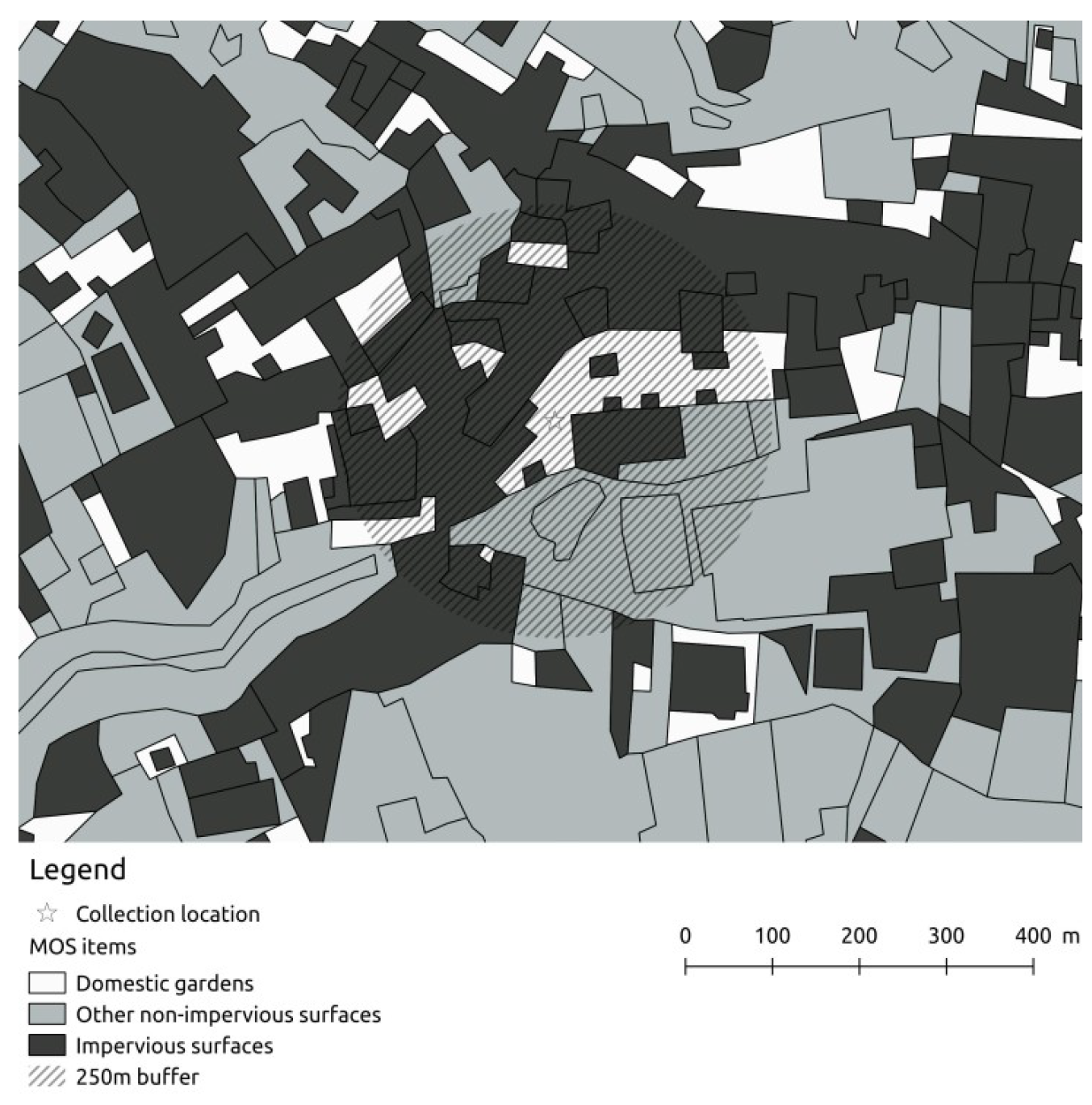
Map extract showing impervious surfaces, domestic gardens, and other non-impervious areas. An example of the SPIPOLL collection location with a 250m buffer.

We computed correlations between the proportion of impervious surfaces, the proportion of domestic gardens, and the inside or outside garden location (1/0). We used the Spearman correlation method for correlations between the proportion of surface variables and the local context variable.

### 2.3 Regression tree modeling

We used the residuals from the models both standardized and unstandardized for local context to fit the regression trees (see Supplementary Fig. S2 for the validation of residuals). Here, we chose to fit the regression tree models to residuals instead of using an alternative method with spatial autocorrelation to control for confounding variables, because the regression tree allows threshold effects to be modeled. For each buffer and both sets of residuals, we repeated the regression tree construction 500 times, each time using a randomly selected subsample of 2070 data points out of 2470. We then selected the most frequent tree structure and extracted all corresponding regression trees from the pool of 500 (see Supplementary Tables S1 and S2). The variables included in the regression trees were the proportion of impervious and garden areas. Regression trees using the residuals unstandardized for local context also included the local garden context variable (categorical: 0/1). We chose to retain both the domestic garden proportion and the dichotomous local context garden variable, because even if a collection was recorded in close proximity to a domestic garden, the precise location might, for instance, be a street or public space situated nearby. These two collections would have different pieces of information related to the contrasting contexts: for instance, management decisions taken by either an inhabitant or the municipality. To better understand the contribution of all spatial scales to morphotype richness, we repeated the regression tree construction while taking into account the proportion of garden and built-up areas on all scales as well as the garden variable for the tree using residuals unstandardized for local context. To be selected to define a node, a variable had to meet the 0.05 criteria of significance. We used the partykit package from R (Hothorn and Zeileis, 2015; Hothorn et al., 2017).

We calculated the threshold values for nodes, the number of data points included in the terminal nodes, and the associated mean morphotype richness as mean values for all regression trees corresponding to the most frequent one. We compared the mean morphotype richness of nodes in all selected trees using the Mann-Withney-Wilcoxon test to assess the significance in the final tree.

We also retrieved the collections by terminal nodes for spatial scales, which appeared in the trees that took into account all spatial scales. For each node, we calculated the proportion of insects belonging to the different orders and compared the repartition of insect orders between terminal nodes using chi squared tests for the main insect orders in the dataset: i.e. Hymenoptera, Lepidoptera, Diptera, Hemipteran, and Coleoptera.

All statistical analyses were conducted in R (R Development Core Team, 2017).

## 3 Results

### 3.1 Diversity of flower-visiting insect data description

#### 3.1.1. Dataset description

The overall dataset contained 2470 collections recorded by 261 citizen scientists on 18 different flower families. On average, morphotype richness per 20 min was 7.51 +/- 6.12 morphotypes. A summary of the observed insect orders and observed morphotypes per order is given in Table 1. Considering each weather variable independently, collections were most frequently associated with low and irregular wind intensity, 0-25% cloud coverage, or a temperature range of 10-20°C. When all weather variables were combined, the greatest number of collections (17.7%) was made during low wind intensity, with 0-25% cloud coverage, and a temperature range of 20-30°C. Asteraceae flowers were the most sampled family of flowers, being found in 32.1% of collections. A detailed distribution of collections according to flower family is provided in Figure 2.

**Table 1.** Description of the data set according to orders morphotypes belong to, proportion of morphotypes belonging to each order, number of different morphotypes seen in each order and number of morphotypes levels appearing in the data set (over 7 levels).

| Order | % of morphotypes<br>belonging to this order | Number of different<br>morphotypes seen | Number of morphotypes<br>levels appearing |
| --- | --- | --- | --- |
| Blattodea | 0.015 | 1 | 1 |
| Coleoptera | 13.9 | 87 | 5 |
| Dermaptera | 0.078 | 1 | 1 |
| Diptera | 34.9 | 73 | 5 |
| Ephemeroptera | 0.0098 | 1 | 1 |
| Hemiptera | 4.96 | 20 | 4 |
| Hymenoptera | 40.4 | 78 | 6 |
| Lepidoptera | 4.61 | 69 | 5 |
| Mecoptera | 0.14 | 1 | 1 |
| Nevroptera | 0.33 | 2 | 1 |
| Opilion | 0.1 | 1 | 1 |
| Orthoptera | 0.61 | 2 | 2 |

#### 3.1.2. Effects of standardization variables

Both standardized and unstandardized models had similar results. Both included a positive effect of 50-75% cloud coverage in comparison to 0-25% on the log of morphotype richness (β=0.12, p=0.0028 and β=0.12, p=0.0028 in the models standardized and unstandardized for local context, respectively). Temperatures between 20°C and 30°C had a significant positive effect compared to those between 10°C and 20°C (β=0.067, p=0.047 and β=0.067, p=0.047 in the models standardized and unstandardized for local context, respectively). The years 2011 to 2017 had a significant positive effect on richness (βs between 0.26 and 0.50, all p<0.01 and βs between 0.26 and 0.50, all p<0.01 in the models standardized and unstandardized for local context, respectively). Week had a significant positive effect on richness (β=0.87, p<0.0001 and β=0.87, p<0.0001 in the models standardized and unstandardized for local context, respectively), while its quadratic term had a significant negative effect on richness (β=-0.0015, p<0.0001 and β=-0.0015, p<0.0001 in the models standardized and unstandardized for local context, respectively). Latitude, longitude, and other categories of factorial variables did not have a significant effect on morphotype richness. Likewise, the local garden context variable did not have a significant effect compared to locations situated outside gardens in the model standardized for local context.

Several flower families had a significant effect on morphotype richness in comparison to the Asteraceae reference family. However, as this was not the main interest of the study, detailed statistics on the effect of flower families will not be given here. In short, Apiaceae and Araliaceae had a significant positive effect on morphotype richness compared to Asteraceae. Boraginaceae, Caprifoliaceae, Crassulaceae, Fabaceae, Geraniceae, Lamiceae, Papaveraceae, Plantaginaceae, Primulaceae, and Scrophulariaceae, which had a significant negative effect on morphotype richness. Autocorrelation strongly decreased in both GLS model residuals and had only a minimal effect on the first 100m interval (see Supplementary Fig. S3).

### 3.2 Relations between environmental variables

The proportion of impervious surfaces and proportion of domestic gardens were correlated at several spatial scales: 50m (p<0.0001, ρ=0.158), 100m (p<0.0001, ρ=0.0962), and 1000m (p=0.00013, ρ=-0.0769). The proportion of domestic gardens and location inside or outside a garden (1/0) were correlated at the same spatial scales: 50m (p<0.0001, ρ=0.872), 100m (p<0.0001, ρ=0.715), and 1000m (p=0.0011, ρ=-0.0659). The proportion of impervious surfaces and location inside or outside a garden (1/0) were correlated at several spatial scales: 50m (p<0.0001, ρ=0.150), 250m (p<0.0001, ρ=-0.178), 500m (p<0.0001, ρ=-0.294), and 1000m (p<0.0001, ρ=-0.402). Other correlations were not significant. A description of ranges for the proportion of impervious surfaces and garden areas is given in Table 2.

**Table 2.** Description of the variables for the proportion of garden areas and impervious surfaces at all spatial scales.

| Spatial scale : | 50m | 100m | 250m | 500m | 1000m |
| --- | --- | --- | --- | --- | --- |
|  | Proportion of garden areas (%) |  |  |  |  |
| Mean +/- sd | 4.65 +/- 12.4 | 3.90 +/- 8.89 | 0.565 +/- 1.62 | 0.308 +/- 1.06 | 0.178 +/- 0.621 |
| Min | 0 | 0 | 0 | 0 | 0 |
| Max | 76.1 | 43.9 | 27.7 | 18.5 | 15.8 |
|  | Proportion of impervious surfaces (%) |  |  |  |  |
| Mean +/- sd | 40.1 +/- 38.3 | 39.5 +/- 35.3 | 44.8 +/- 31.7 | 41.1 +/- 32.3 | 41.6 +/- 31.2 |
| Min | 0 | 0 | 0 | 0 | 0 |
| Max | 100 | 100 | 100 | 100 | 98.2 |

### 3.3 Selected trees after bootstrapping

Selected trees were represented by 42.2% to 82.4% of the pool of 500 regression trees depending on the buffer scale and the model standardized or unstandardized for local context. Details of all regression tree structures observed in the pool of 500 trees are provided in Supplementary Tables S1 and S2 for the models standardized and unstandardized for local context, respectively.

Tree structures resulted from only one division, leading to trees with two nodes, or from two divisions, a first-order division leading to two branches and a second-order division in one of the branches, leading to a final number of three nodes.

### 3.4 Role of spatial scales and environmental variables in tree structures

In the regression trees using residuals from the model unstandardized for local context, the consideration of all spatial scales resulted in the proportion of garden areas in the 50m buffer being the first structuring variable at a threshold of 29.3 +/- 2.8% (Fig. 4). When focusing on the tree using variables of the 50m buffer only, this variable remained the first structuring variable at a threshold of 29.4 +/- 3.0% (Fig. 5A). Regression trees for 100m and higher buffers all resulted in the local garden context variable (0/1) as a structuring variable. At the buffer spatial scale of 100m (Fig. 5B), the garden context (0/1) was the first structuring variable, while the proportion of garden area was the second-order division. At the buffer spatial scales of 250m and 1000m (Fig. 5C, E), the garden context (0/1) was the first structuring variable, while the proportion of impervious surface was the second-order division in the garden context 1 branch. This was reversed at the buffer spatial scale of 500m (Fig. 5D).

**Figure 4.**
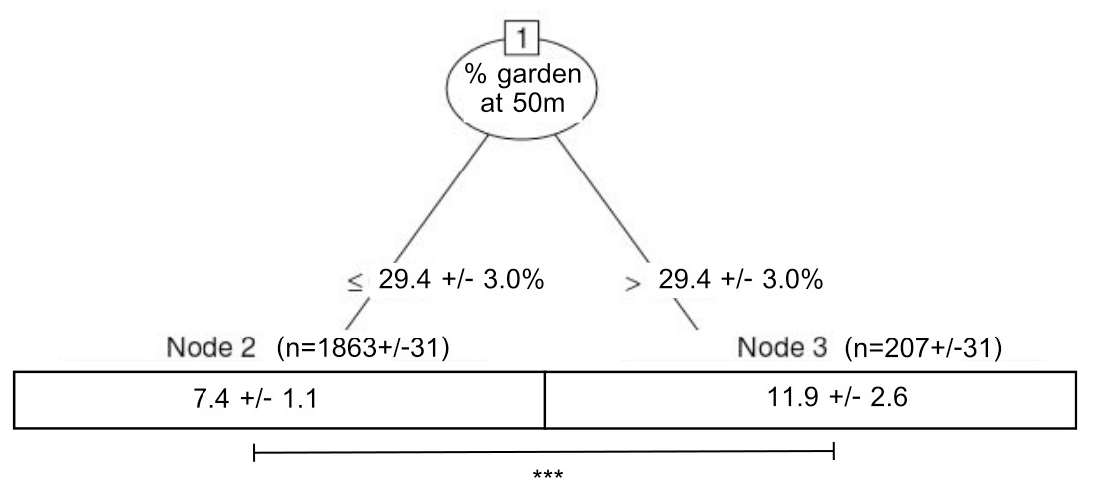
Regression tree for the model unstandardized for local context (i.e., excluding the garden context variable) with all spatial scales taken into account and a synthesis of all corresponding trees (n=244) from the pool of 500 drawn trees. Variables for nodes are shown in circles. Threshold values of the variables are given on the formed branches. Terminal nodes are numbered, and the associated mean +/- sd for morphotype richness is stated. Significant p-values for the difference between mean morphotype richness are indicated.

**Figure 5.**
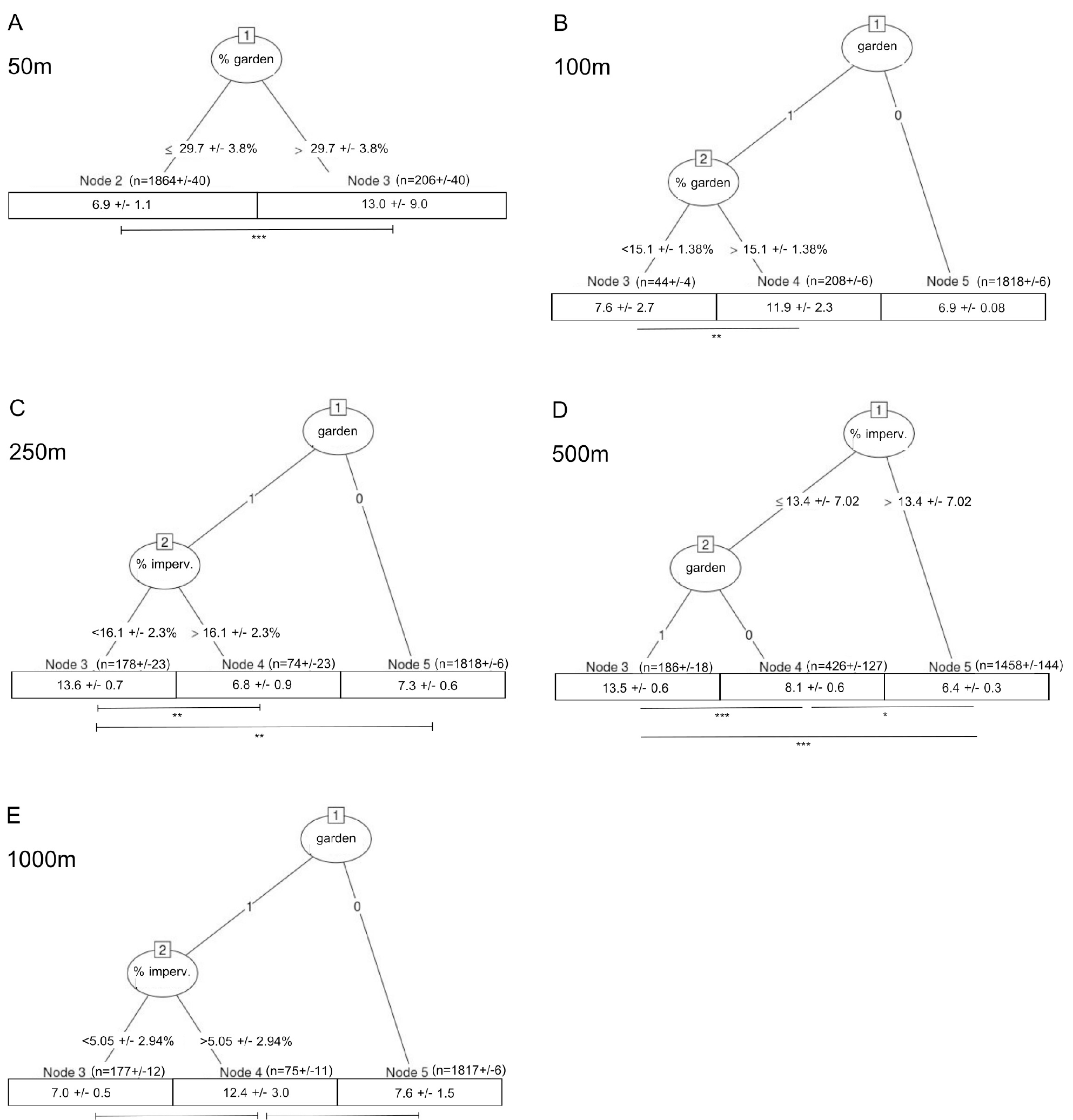
Regression trees for the model unstandardized for local context (i.e., excluding the garden context variable). A: 50m buffer; B: 100m buffer; C: 250m buffer; D: 500m buffer; E: 1000m buffer. Variables for nodes are shown in circles with the associated p-value. Threshold values of the variables are given on the formed branches. Terminal nodes are numbered, and the associated mean +/- sd for morphotype richness is provided. Significant p-values for the difference between mean morphotype richness are indicated below each tree.

As the local garden context variable appeared in all trees except two when using residuals from the model unstandardized for local context, it may have masked the effect of the proportion of garden area variable. Regression trees using residuals from the model standardized for local context helped disentangle this.

In the regression trees using residuals from the model standardized for local context, taking all spatial scales into account resulted in a tree similar to when the local garden variable was included: the proportion of garden area in the 50m buffer was the first and only structuring variable at a threshold of 29.7 +/- 3.8%. At a spatial scale of 100m, the proportion of garden area was also the first structuring variable at a threshold of 22.6 +/- 2.24%. The branch with a low garden proportion was further divided, again with the proportion of garden areas as a structuring variable at a threshold of 6.38 +/- 1.43%. At spatial scales of 250m and higher, the proportion of impervious surface was the first structuring variable. It was also the only structuring variable except at the 1000m spatial scale, which involved the proportion of garden areas in the low impervious surface branch in a second-order division. Regression trees using residuals from the model standardized for local context can be found in Supplementary Figure S4.

Table 3 summarizes the steps in the regression trees at which the proportion of the garden area, the local garden context variable, and the proportion of impervious surfaces had an effect.

**Table 3.** Summary of variable effects in the regression trees at the different spatial scales. Mention of 1 or 2 indicates if the variable was used in order to define a first order division in the tree of a second order division, i.e. inside an already defined branch of the tree. Results for trees without the local garden context variable are shown in brackets.

|  | Local | → | → | → | Landscape |
| --- | --- | --- | --- | --- | --- |
| Spatial scale | 50m | 100m | 250m | 500m | 1000m |
| Context | - | 1 (-) | 1 (-) | 2 (-) | 1 (-) |
| Garden area | 1 (1) | 2 (1, 2) | - | - | - (2) |
| Impervious surface | - | - | 2 (1) | 1 (1) | 2 (1, 2) |

### 3.5 Dependence of pollinator richness on spatial scales and environmental variables

Maximal morphotype richness was reached in nodes attached to the branch with a higher proportion of garden areas or a lower proportion of impervious surfaces in the two-node trees (Fig. 5, Supplementary Fig. S4). In the three-node trees, we observed that a node with maximal morphotype richness could not always be identified at a 0.05 significance level. However, such a node could be identified in trees with a spatial scale of 250m, 500m, and 1000m using residuals from the model unstandardized for local context (Fig. 5) and in trees with a spatial scale of 100m and 500m using residuals from the model standardized for local context. In all but one of the five concerned trees, maximal morphotype richness was reached in a node resulting from a low proportion of impervious surfaces combined with either a high proportion of garden areas or a location inside a garden (local garden context variable = 1). In the last tree, using residuals from the model unstandardized for local context at a spatial scale of 500m (Fig. 5E), maximal morphotype richness was reached in a node resulting from a combination of location inside a garden and a high proportion of impervious surfaces.

### 3.6 Repartition of orders between terminal nodes in trees with the domestic garden proportion variable

The proportion of domestic gardens was a significant explanatory variable in regression trees taking into account all spatial scales and using residuals from both models. It was significant at the 50m spatial scale only in these regression trees, which had two terminal nodes according to a lower or higher proportion of domestic gardens at 50m.

In both regression trees, the patterns of response of insect orders were similar. The terminal node determined by a proportion of domestic garden less than 29.4% in the tree based on the unstandardized model (compared to 29.7% in the tree based on the standardized model) encompassed 2217 collections (compared to 2218 in the tree based on the standardized model), while the proportions of insects belonging to Lepidoptera (chi^2^=23.85, p<0.001 and chi^2^=23.4, p<0.001 for the unstandardized and standardized models, respectively) and Diptera (chi^2^=20.9, p<0.001 and chi^2^=21.9, p<0.001 for the unstandardized and the standardized models, respectively) were lower in this node compared to the node determined by a proportion of domestic gardens exceeding 29.4% (29.7% in the standardized model). The low domestic garden proportion node also encompassed a higher proportion of insects belonging to Coleoptera (chi^2^=53.2, p<0.001 and chi^2^=52.7, p<0.001 for the unstandardized and the standardized models, respectively) compared to the high domestic garden proportion node.

## 4 Discussion

The present study considered impervious surfaces and domestic garden areas together in order to understand their relative effects on pollinator richness. As domestic gardens are difficult to access and study, but nevertheless important places where various and numerous pro-biodiversity actions may take place, understanding their role is relevant. By introducing a garden context variable, we were also able to retain information about the local context of data points. Indeed, domestic garden proportions were not always correlated to the garden context variable, and thus both variables retained information about the situation of data points relative to domestic gardens.

### 4.1 Local and landscape determinants of pollinator richness

Maximal morphotype richness of pollinators was associated with data points that were located inside gardens, surrounded by a higher proportion of gardens, surrounded by a lower proportion of impervious surfaces, or a combination of these criteria. This is consistent with previous studies investigating the effect of urbanization on pollinators: urban areas have negative effects on both wild bee (Geslin et al., 2016) and moth richness (Bates et al., 2014) and more generally limit plant-pollinator interactions (Desaegher et al., 2017). The mean morphotype richness of pollinators found in this study – 7.51 morphotypes per 20 minutes – is also consistent with a previous measure based on the SPIPOLL database and focusing on eight plant species: 4.47 morphotypes per 20 minutes (Deguines et al., 2016).

We did not include finer scales of pollinator diversity in the regression tree analysis, as total richness is an appropriate surrogate for diversity when dealing with citizen science data (Kremen et al., 2011). The consistency of our results at different spatial scales nevertheless indicates a general pattern of pollinator richness in the densely urbanized Île-de-France region, even if responses of individual species may vary.

In spite of its high levels of urbanization, Île-de-France contains forests and farming areas as well as semi-natural areas and other UGS. Farming areas usually have less pollinator richness than urban residential areas (Baldock et al., 2015), while urban parks, natural areas, and urban remnants have more pollinator richness than residential and domestic garden areas (Hostetler and McIntyre, 2001; Matteson et al., 2013; Tommasi et al., 2004). We did not directly compare the impact of domestic garden areas with that of other permeable land use areas known to favor pollinators, but our results still provided insights into these variations: the high morphotype richness found in data points inside gardens and within higher impervious surface proportion at the 1000m scale may indicate that non-impervious surfaces are not all equivalent. We may here point to the role of farming areas. At the 1000m scale, farming areas, which are negatively related to pollinator richness (Baldock et al., 2015), may have been largely included in the buffers. Indeed, the Île-de-France region, apart from dense urbanization, includes large areas of conventional wheat farming. A negative effect of these farming areas might be related to the observed higher richness inside gardens and within areas of higher impervious surfaces, i.e. away from farming areas. Such data points with high morphotype richness inside gardens and within areas of higher impervious surfaces at the 1000m scale are not numerous (n=75 +/- 11 depending on the bootstrapped tree). They may also represent a few small areas of good habitat within a large matrix of non-favorable habitat, with the mean richness reflecting the situation of a single almost inaccessible favorable patch (Steffan-Dewenter et al., 2002).

### 4.2 Domestic gardens as favorable habitats within less favorable impervious surfaces

Garden area effects, i.e. effects of being surrounded by domestic gardens, were only seen at smaller scales, while impervious surface effects, i.e. effects of being surrounded by impervious surfaces, were at larger scales. The garden variable, always representing a local effect, appeared to be a strong structuring variable at almost all spatial scales. This variable conveys a complementary significance regarding the proportion of domestic gardens, as both variables are not correlated at the intermediate spatial scale, i.e. 250m and 500m. Being located in a domestic garden seems to be always favorable at nearly all spatial scales, while being surrounded by domestic gardens exerted a positive effect only at small spatial scales, i.e. 50m and 100m. Impervious surfaces seem to act as strong determinants for pollinator richness, likely determining the richness found at a given place according to pollinator affinity for impervious and urban areas (Deguines et al., 2012). At large scales, a patch of favorable local habitat may not be reached if it is surrounded by an unfavorable landscape, as shown for butterflies (Bergerot et al., 2011). Once a determined diversity of pollinators is hosted by a region encompassing impervious surfaces, gardens are more likely to host a greater proportion of this diversity compared to other areas in the same region.

Pollinator species perceive landscape structures at different spatial scales depending on their dispersion or flight ranges (Keitt et al., 1997; Steffan-Dewenter et al., 2002; With et al., 1999). Analyzing several Apidae groups relative to the features of numerous spatial scales, Steffan-Dewenter et al. (2002) found solitary bee richness to be positively correlated to the proportion of semi-natural areas up to 750m, contrary to honeybee and bumblebee richness, with this pattern being reversed over 750m. Here, the 50m spatial scale appeared to be significant when all spatial scales were analyzed together, while changes in the dominant structuring variables occurred at a 250m spatial scale when spatial scales were analyzed separately. The effects of local and landscape levels are likely to rely on different species responses: local features, which disperse in larger scales, may be quite important for some species.

The type of identification used in the SPIPOLL program does not allow this finer taxonomic scale of pollinator diversity to be studied. It also results in a lower diversity of observable taxonomic groups, which may have consequently led to lower variability in the data to some extent. This may be viewed as positive, as we were nevertheless able to show significant and consistent patterns. While citizen scientist protocols have been shown to overestimate rare species and underestimate common ones, the amount of data collected is likely to compensate for this trend (Gardiner et al., 2012), and the use of a computer-aided identification tool, here the SPIPOLL online identification key, ensures reliable identification and thus consistent sorting of taxa across citizen scientists and stable morphotype categories (Obrist and Duelli, 2010). The measures of morphotype richness taken by citizen scientists have been previously validated for the detection of changes in abundance, richness, or similarity over space and time (Kremen et al., 2011), allowing for ecological patterns to be identified (Hochachka et al., 2012) and SPIPOLL citizen scientists to improve their knowledge about pollinator identification while participating (Deguines et al., 2018). Morphotypes themselves are relevant for the calculation of ecological indicators, such as pollinator diversity (Obrist and Duelli, 2010). Moreover, the citizen science program has economic advantages, as the field work needed in order to collect an equivalent pool of data would have required a large amount of money; indeed, the SPIPOLL program has much lower costs, with estimated annual savings between 678,523€ and 4,415,251€ in France (Levrel et al., 2010). Our use of the SPIPOLL citizen science database allowed us to gather a large amount of data (n=2470 data points), including inside domestic gardens, which are less easily accessible (Hernandez et al., 2009).

### 4.3 Resulting thresholds and difficulties arising from their use

Regression trees allowed us to identify threshold effects and cascading thresholds as opposed to linear effects. At small scales, the identified thresholds are likely to represent typical situations of residential housing. For instance, the threshold for the proportion of garden areas is 29.7 +/- 3.8% at 50m and 15.7% at 100m, which may account for locations situated between gardens, houses, and access roads. At small spatial scales, impervious surfaces and domestic gardens surfaces proportions were correlated and impervious surfaces co-occurred with gardens location. This is likely to indicate this parallel evolution of housing with dwellings and gardens on the one hand, and residential infrastructure with access roads and other impervious surfaces on the other. The threshold figures might seem smaller than what could be expected for the respective proportions of detached houses, gardens, and access roads. This seemingly small figure may be compared to the results pertaining to population conservation, which requires larger areas than those identified for individual species conservation (Beninde et al., 2015; Smith, 2007). The identified threshold here is only a minimum corresponding to the species observed by participants, which is a subset of total diversity.

Considering an average garden size of 571m^2^ (Dewaelheyns et al., 2014), the proportions of domestic gardens in both 50m and 100m buffers represent approximately four gardens in the 50m buffer and eight gardens in the 100m buffer. Assembled in a patch, these gardens may account for coordinated individual actions in favor of pollinators in the neighboring gardens, either separated by fences or situated in a row along a street. As a result, it would not require many inhabitants to transform individual decisions with uncertain collective consequences into coordinated gardening decisions in favor of the target organism, that is, to shift from the “tyranny of small decisions” to a “resource by small gardening actions” (Dewaelheyns et al., 2016).

The threshold for the proportion of built-up areas with an effect on morphotype richness is variable in regression trees, ranging from 250m to 1000m scales. Considering the variety of pollinators, it may be that some key morphotypes respond overwhelmingly to certain spatial scales at different thresholds, which would result in these inconsistencies.

We chose to use the proportion of garden and built-up areas as landscape indicators in this study. Correlations exist between both proportions, but are not present at all spatial scales and are of low value when present, i.e. less than 0.16. Thus, each indicator independently contributes information in this study. A similar surface proportion of the indicator, regardless of which one, may point to a situation with either numerous fragmented areas or a single large area: total surface and fragmentation degree are thus undifferentiated (Andrén, 1994). Here, we cannot assess the particular situation to which pollinator richness responded. Although threshold determination is useful in statistical modeling when the non-linear effect is hypothesized, thresholds should subsequently be manipulated with caution. To guide urban planning in favor of biodiversity, goals and limitations associated with these thresholds need to be explicit from both scientific and management perspectives (Beninde et al., 2015). Indeed, each threshold may be viewed from two angles: either as a minimal requirement to be expanded or as a limiting value to be reached.

### 4.4 Orders favored at smaller scales in gardens

Finer scales of pollinator diversity may reveal a variability of responses to urbanization, as some groups, such as cavity-nesting bees (Hernandez et al., 2009) and social small-bodied species (Banaszak-Cibicka and Zmihorski, 2012), increase in cities, while others decrease: pollinators thus have various levels of urbanophobia (Deguines et al., 2012). Here, we identified orders that were favored or not by domestic garden surroundings at small spatial scales. Lepidoptera were favored by domestic garden surroundings at the 50m spatial scale, which is consistent with their known sensibility to urbanization (Ramirez-Restrepo et al., 2017) and negative affinity for urban land use (Deguines et al., 2012). Likewise, Diptera have rather low affinities for urban land use (Deguines et al., 2012), which is consistent with them being favored by domestic garden surroundings in this study. Hymenoptera were not favored by domestic garden surroundings at the 50m spatial scale. As insects belonging to this order are likely to have a positive affinity for urban land use (Deguines et al., 2012), the proportion of domestic gardens taken into account in this study may not be sufficient to reveal a positive effect of these less impervious surfaces (McKinney, 2008). Variability of species responses within the order may also account for the absence of differences when taking into account the entire order. Here, we also revealed that Coleoptera are not favored by domestic garden surroundings, which may be surprising given the general negative effect of urban land use on insects (McKinney, 2008; McIntyre, 2000), but this may also reflect certain management practices in gardens.

Overall, our findings indicate that Lepidoptera and Diptera could benefit from combined decisions in favor of insects at small spatial scales, coordinated between four and eight domestic gardens. This also points to the necessity for broader communication about insects and pollinators, as they are usually associated with negative feelings (Kellert, 1993), but can be recognized for utilitarian or aesthetic values (Barrow, 2002). While Lepidoptera have aesthetic qualities (Barrow, 2002), this is not the case with Diptera, for which more knowledge diffusion could help garden management in their favor.

## 5 Conclusion

This study identified the less explored role of domestic gardens as places that structure favorable features for pollinator habitat selection at the local scale after filtering due to urbanization. In line with Goddard et al. (2010), we acknowledge that the consideration of garden owners’ individual management decisions as a whole is problematic as the scale of management is, at least, order-dependent. Garden owners should be encouraged to focus on management favorable to species with short dispersion and flight ranges, as these are most likely to be beneficial given that the positive effect of gardens occurs at this spatial scale. Concerning specific orders, garden owners should be encouraged to focus on management favorable to Lepidoptera and given more information about Diptera. To outline propositions for conservation behavior at the domestic garden scale, our study could be repeated with a smaller taxonomic resolution. For example, it could be extended to unique morphospecies. Public policies will be more effective if they combine various management approaches based on features that impact pollinator richness at different scales and favor connectivity instead of isolated patches, as well as coordination between local actors, namely gardeners, at the scale of garden patches. Regarding the SPIPOLL program, using morphotype identification level, which is less precise than species identification level, is also linked to the acknowledgment of participants’ work. Indeed, using this dataset and communicating the results to participants is a means to motivate citizen scientists to continue contributing to the program.

## Acknowledgments

The authors acknowledge all SPIPOLL participants, the SPIPOLL database managers, as well as two anonymous reviewers who provided helpful comments.

## Funding

Île-de-France PSDR Dynamiques project, Labex BASC. M.L. is funded by the French Ministry of Research and the FdV Doctoral School.

